# Inhibition of the androgen-activating enzyme AKR1C3 selectively decreases systemic and intra-adipose 11-oxygenated androgens in women

**DOI:** 10.64898/2026.03.27.714735

**Authors:** Lina Schiffer, Amarah V. Anthony, Laura B. L. Wittemans, Angela E. Taylor, Imken Oestlund, Antonio M.A. Miranda, Eka Melson, Tara McDonnell, Punith Kempegowda, Paul Smith, T. Justin Clark, Martin Wabitsch, Michael W. O’Reilly, Michaele Peters, Andrea Wagenfeld, Jan-Peter Ingwersen, Jacky L. Snoep, William R. Scott, Jan Hilpert, Karl-Heinz Storbeck, Wiebke Arlt

**Affiliations:** MRC Laboratory of Medical Sciences, London, W12 0HS, UK; Big Data Institute at the Li Ka Shing Centre for Health Information and Discovery, University of Oxford; Oxford, OX3 7LF, UK; Nuffield Department of Women’s and Reproductive Health, Medical Sciences Division, University of Oxford, Oxford, OX3 9DU, UK; Novo Nordisk Research Centre Oxford, Oxford, OX3 7FZ, UK; Department of Metabolism and Systems Sciences, School of Medical Sciences, College of Medical and Health Sciences, University of Birmingham, Birmingham, B15 2TT, UK; Department of Biochemistry, Stellenbosch University, Stellenbosch, 7602, South Africa; Institute of Clinical Sciences, Faculty of Medicine, Imperial College London, London, W12 0NN, UK; Department of Medicine, Royal College of Surgeons in Ireland, University of Medicine and Health Sciences, Dublin, D09 V2N0, Ireland; Department of Applied Health Sciences, School of Health Sciences, College of Medical and Health Sciences, University of Birmingham, Birmingham, B15 2TT, UK; Birmingham Women’s Hospital, Birmingham Women’s and Children’s NHS Foundation Trust, Birmingham, B15 2TG, UK; German Center for Child and Adolescent Health (DZKJ; partner site Ulm), Division of Pediatric Endocrinology and Diabetes, University Hospital of Ulm, Ulm, 89075, Germany; Research and Development, Pharmaceuticals, Bayer AG, Berlin, 13353, Germany; Vrije Universiteit Amsterdam, Systems Biology Lab, AIMMS/A-LIFE, Amsterdam 1081, The Netherlands

## Abstract

Androgen excess drives metabolic and reproductive complications in polycystic ovary syndrome (PCOS), affecting 10-15% of women globally. Aldo-keto reductase 1C3 (AKR1C3) converts inactive precursors from both the classic and the recently identified 11-oxygenated androgen pathways, generating testosterone and 11-ketotestosterone, respectively, which exert comparable androgen receptor activation. Both circulate in similar concentrations in premenopausal women while 11-ketotestosterone is predominant after menopause and in PCOS. Here, we show that adipocytes are a major site of *AKR1C3* and androgen receptor expression, with increased expression in women and individuals with obesity. Using human female adipose tissue explants, we find a much higher activation of 11-oxygenated over classic androgens, observing a decrease in 11-oxygenated but not classic androgen activation by AKR1C3 inhibition. Correspondingly, we demonstrate that AKR1C3 inhibitor treatment in premenopausal women selectively disrupts the activation of 11-oxygenated androgens. Pharmacological targeting of AKR1C3 provides a novel strategy to alleviate systemic and intra-adipose 11-oxygenated androgen excess.

**One Sentence Summary:** Inhibition of the androgen-activating enzyme AKR1C3 results in a major decrease in 11-oxygenated but not classic androgens in women.

---

Polycystic ovary syndrome (PCOS) is the most common endocrine disorder in women, with a global prevalence of 10-13% ^1^. PCOS manifests with an increased life-long risk for the development of metabolic disease ^1^, creating major burdens to affected women and healthcare systems ^2^. Androgen excess is a defining feature of PCOS and has been implicated as a major driver of the associated life-long metabolic complications ^1,3^, including insulin resistance, dysglycemia, type 2 diabetes mellitus ^4–8^, and metabolic dysfunction-associated steatotic liver disease (MASLD) ^9–11^. Despite its high worldwide prevalence, there is currently no pharmacological therapy approved for specific use in women with PCOS and there is no known specific intervention to prevent and treat androgen excess-induced metabolic dysfunction.

We have previously demonstrated that adipose tissue is a key metabolic compartment linking androgen action and systemic metabolic dysfunction in women with PCOS ^12,13^. Intra-adipose conversion of the androgen precursor androstenedione (A4) to the potent classic androgen testosterone (T), catalyzed by the enzyme aldo-keto reductase 1C3 (AKR1C3), drives lipid accumulation in adipocytes and promotes a systemic lipotoxic metabolome ^12,14^, which in turn results in hyperinsulinemia. Notably, *AKR1C3* expression in adipocytes is upregulated by insulin ^12,15^, thereby creating a vicious circle linking enhanced androgen activation by AKR1C3 with increased insulin resistance.

Recent research has uncovered the existence of a new androgen class, the 11-oxygenated androgens, which are derived from the peripheral metabolism of the abundant adrenal-derived steroid 11β-hydroxyandrostenedione (11OHA4) ^16–18^. 11OHA4 is converted to 11-ketoandrostenedione (11KA4), which is then activated to 11-ketotestosterone (11KT) by AKR1C3, which also converts A4 to T (**Fig. 1A**). 11KT activates the androgen receptor with similar potency and efficacy as T ^19,20^, and circulates at concentrations equal to or higher than testosterone in women ^21,22^. However, in contrast to T, 11KT does not decline with age in women ^21–23^ and, therefore, represents the dominant androgen after menopause, a period characterized by a significant increase in metabolic dysfunction ^24^. 11-oxygenated androgens have been shown to be increased in women with PCOS where they represent a major fraction of circulating androgens ^25,26^. Several studies in women with PCOS reported correlations between 11-oxygenated androgens and markers of insulin resistance ^25,26^, though a differential role of 11-oxygenated androgens over classic androgens in promoting metabolic dysfunction in PCOS has not been delineated yet.

**Fig. 1.**
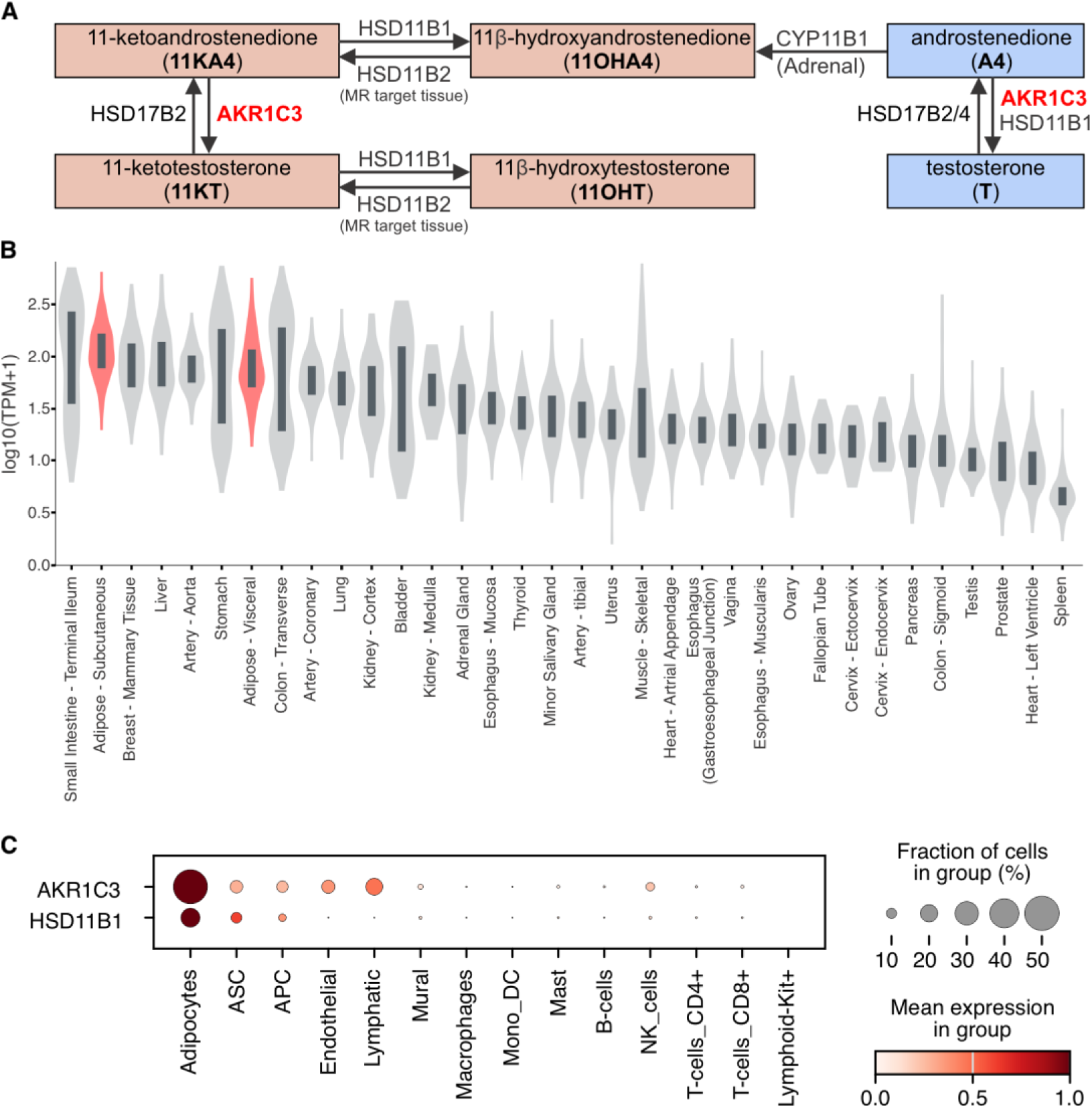
Expression of *AKR1C3* in adipose tissue. (**A**) Schematic showing the roles of AKR1C3, HSD11B1 and other enzymes in classic and 11-oxygenated androgen biosynthesis. (**B**) Bulk RNAseq data from the Adult Genotype-Tissue Expression (GTEx) Project (dbGaP Accession phs000424.v10.p2) showing that subcutaneous and visceral adipose (red) are among the tissues with the highest median *AKR1C3* expression. The dark grey boxes indicate the 25^th^ to 75^th^ percentile, while the violin plots indicate the distribution density. (**C**) Dot plot (mean expression scaled) from snRNA-Seq of human subcutaneous adipose tissue (n=49) showing that *AKR1C3* and *HSD11B1* expression is most abundant in adipocytes (ASC, adipose-derived stem cells; APC, adipose progenitor cells; Mono_DC, monocyte-derived dendritic cells; NK_cells, natural killer cells).

Here, we examined the differential contributions of the classic and the 11-oxygenated androgen pathways to androgen biosynthesis in female adipose tissue and the impact of selective inhibition of AKR1C3 on systemic and adipose-specific androgen production. We did so to explore a potential therapeutic role of AKR1C3 inhibitors in treating androgen excess and metabolic dysfunction in women with PCOS.

## RESULTS

### Adipocytes are the major site of *AKR1C3* expression in adipose tissue

Analysis of bulk RNA-Seq data from the Adult Genotype Tissue Expression Project (GTEx) revealed that adipose tissue ranks amongst the tissues with the highest *AKR1C3* expression, with higher expression in subcutaneous than visceral adipose tissue (**Fig. 1B**).

To investigate which specific cell types within adipose tissue express *AKR1C3*, we interrogated a single nucleus RNA-Seq (snRNA-Seq) dataset derived from adipose tissue biopsies obtained from male and female individuals with obesity (n=34; 25 female) and healthy lean controls (n=28; 18 female) ^27^. This analysis established adipocytes as the cell type with predominant *AKR1C3* expression in adipose tissue (**Fig. 1C**). Importantly, this expression pattern across cell types, i.e. the highest expression in adipocytes, was mirrored by androgen receptor (*AR*) expression (**Extended Data Fig. 1A**).

The enzyme HSD11B1, which inactivates 11-oxygenated androgens ^28^ but activates classic androgens ^29^ (**Fig. 1A**), also showed highest expression in adipocytes, followed by precursor cells, i.e. adipose-derived stem cells (ASC) and adipose progenitor cells (APC), but with minimal expression in other cell types (**Fig. 1C**).

### Sex-specific differences in adipose *AKR1C3* expression and relationship to BMI and obesity

Analysis of the bulk RNA-Seq GTEx data showed that *AKR1C3* expression is significantly higher in women than in men in subcutaneous (**Fig 2A**, p=2.34e-05) but not in visceral (**Fig. 2B**) adipose tissue. Analysis of the bulk RNA-Seq data also revealed that *AKR1C3* expression in subcutaneous adipose tissue positively correlated with BMI in women (R=0.23, p=0.0063) but not in men (p=0.04) (**Fig. 2A**). By contrast, *AKR1C3* expression in visceral adipose tissue correlated with BMI in both sexes (**Fig. 2B**, R=0.21, p=0.017 for female; R=0.17, p=0.0029 for male adipose tissue).

**Fig. 2.**
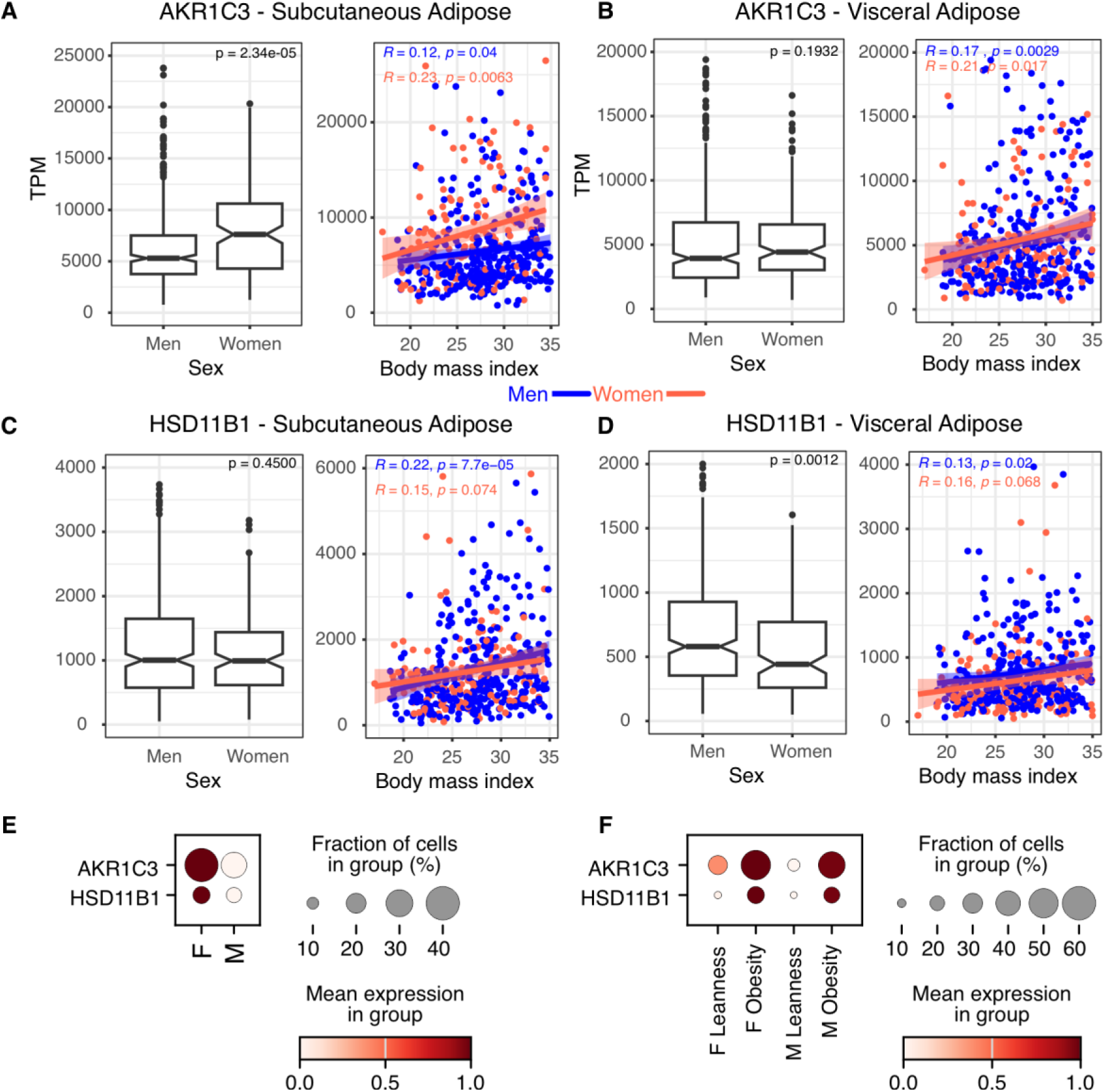
The influence of sex and BMI on *AKR1C3* and *HSD11B1* expression in adipose tissue. Analysis of bulk RNA-Seq data from the Adult Genotype-Tissue Expression (GTEx) Project (dbGaP Accession phs000424.v10.p2) showing the relationship of sex (Tukey box-and-whisker plots, p-values determined by Mann-Whitney U test) and BMI (Pearson correlation with 95% confidence interval) with *AKR1C3* and *HSD11B1* expression in subcutaneous (**A & C**) and visceral adipose tissue (**B & D**). (**E**) Dot plot (mean expression scaled) from snRNA-Seq analysis of subcutaneous adipocytes from female (F, n=33) and male (M, n=16) donors showing the differential expression of *AKR1C3* and *HSD11B1*. (**F**) Dot plot (mean expression scaled) from snRNA-Seq analysis comparing *AKR1C3* and *HSD11B1* expression in subcutaneous adipose tissue from 24 individuals with leanness (15 female) and 25 with obesity (18 female).

In the GTEx data, *HSD11B1* expression in subcutaneous adipose was lower than *AKR1C3* expression, did not differ between sexes in subcutaneous adipose (p=0.45) (**Fig 2C**), but was significantly higher in visceral adipose tissue from men compared to women (**Fig 2D**, p=0.0012). *HSD11B1* expression in adipose tissue correlated positively with BMI in men (subcutaneous: R=0.22, p=7.7e-5; visceral: R=0.13, p=0.02), but not in women (p>0.05) (**Fig. 2C and D**). Analysis of the snRNA-Seq data showed that expression levels of *AKR1C3* and *HSD11B1* in subcutaneous adipocytes were higher in women than men (p= 1.76e-39 for AKR1C3 and p= 0.008 for HSD11B1, **Fig. 2E and Extended Data Table 1**), again with higher expression of *AKR1C3* than *HSD11B1*. Donors with obesity had higher mRNA levels of *AKR1C3* and *HSD11B1* than lean donors in both sexes (all p<1e-23; **Fig. 2F and Extended Data Table 1**).

In the bulk RNA-Seq analysis, expression of *AR* showed no sex differences or correlation with BMI of subcutaneous adipose tissue (p=0.23, **Extended Data Fig. 1B**). However, analysis of the snRNA-Seq data revealed higher AR expression in subcutaneous adipocytes from women compared to men (p=6.71e-9; **Extended Data Fig. 1C**) and from women with obesity compared to lean women (p=8.22e-148; **Extended Data Fig. 1D**). In visceral adipose tissue, *AR* expression was higher in women than men (p=5.93e-05) and correlated negatively with BMI in both men (R=-0.18, p=0.0017) and women (R=-0.28, p=0.0013; **Extended Data Fig. 1E**).

### Human female adipose tissue preferentially activates 11-oxygenated androgens

Next, we determined AKR1C3-mediated activation of classic and 11-oxygenated androgens in *ex vivo* experiments using human female adipose tissue. We collected subcutaneous and visceral adipose tissue from women during elective surgery and in a paired tissue design incubated them with the AKR1C3 substrates androstenedione (A4; classic androgen precursor) and 11KA4 (11-oxygenated androgen precursor; see **Fig. 1A**). To cover a broad BMI range, we included both women undergoing elective bariatric surgery (n=8; age range 32-59 years, BMI range 42.0-56.8 kg m^-2^) and women undergoing elective abdominal surgery for non-cancerous, gynecological indications (n=22; age range 36-74 years, BMI range 20.8-38.2 kg m^-2^). In both subcutaneous and visceral adipose tissue, we observed predominant activation of 11-oxygenated androgens, i.e. conversion of 11KA4 to 11KT and its immediate downstream metabolite 11β-hydroxytestosterone (11OHT), a partial androgen receptor agonist ^19,20^, with only small amounts of A4 activated to T in the classic androgen pathway. 11KT generation was more than 10-fold higher than that of T in subcutaneous (12.4-fold) and visceral (10.5-fold) female adipose tissue (**Fig. 3A and B; Extended Data Fig. 2A-D**).

**Fig. 3.**
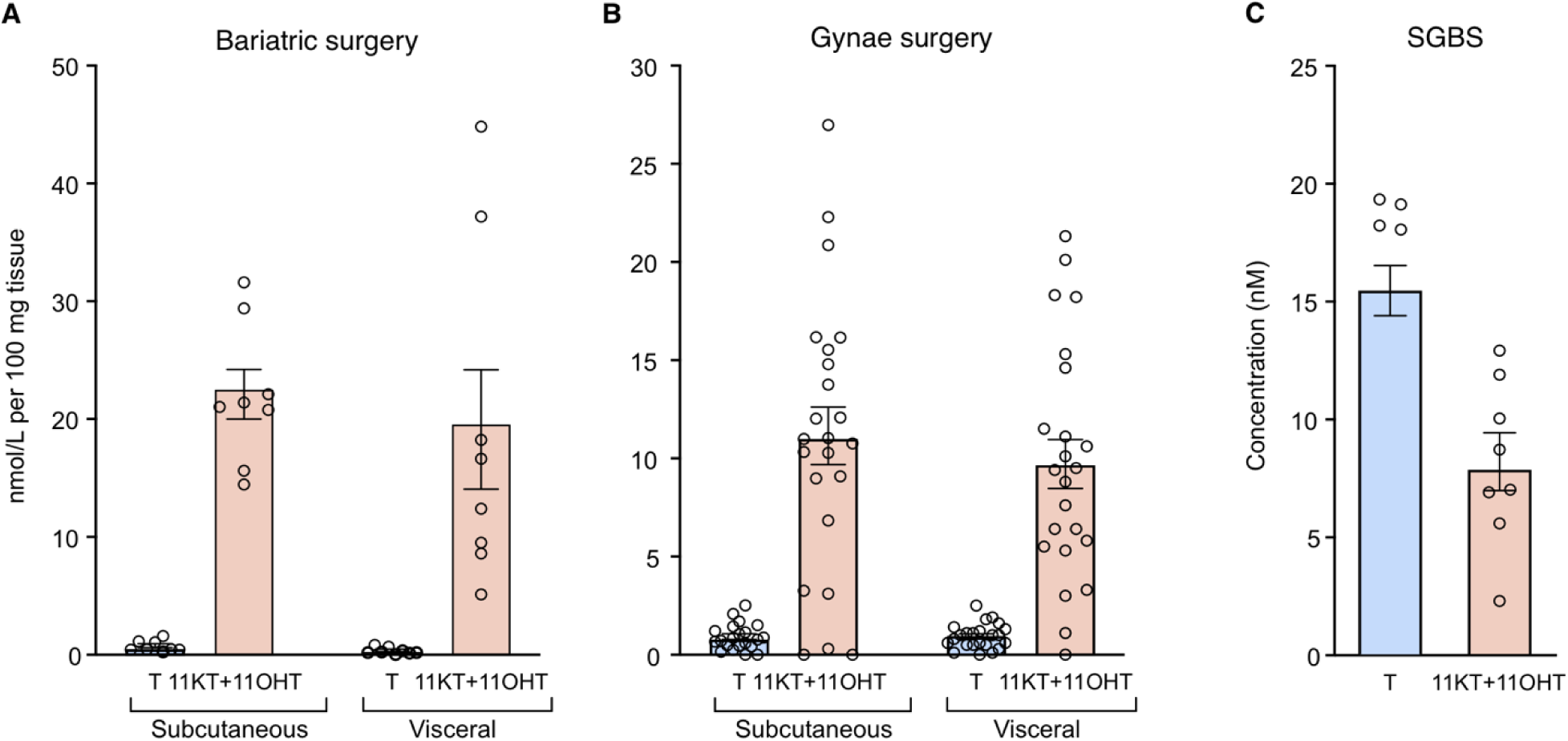
*Ex vivo* metabolism of classic and 11-oxgenated androgens in human female adipose tissue. Incubations of subcutaneous and visceral adipose tissue showed preferential generation of active 11-oxygenated androgens (11KT and 11OHT) over the classic androgen (T). Adipose tissue was collected from women undergoing (**A**) bariatric surgery (n=8, age 32-59 yrs, BMI 42.0-56.8 kg/m^2^) and (**B**) gynecological surgery (n=22, age 36-63 yrs, BMI 20.8-38.2 kg/m^2^) and incubated with the AKR1C3 substrates androstenedione or 11KA4 (100 nM) for 72h (bariatric surgery) or 24h (gynecological surgery). Androgen metabolites were quantified by LC-MS/MS. (**C**) Incubation of differentiated SGBS adipocytes with A4 or 11KA4 (100 nM) results in greater generation of the active classic androgen (T) compared to the 11-oxygenated androgens (11KT and 11OHT) (n=8 or 9). Bars show the mean of all independent experiments ± SEM; circles represent the individual tissue donors (A and B) or independent experiments (C).

We also used differentiated Simpson-Golabi-Behmel Syndrome (SGBS) pre-adipocytes for incubations with classic and 11-oxygenated androgen precursors, as SGBS cells are a commonly used *in vitro* model to study human adipose biology including hormonal dysregulation in PCOS ^14,15,30^. SGBS cells are derived from fibroblasts obtained post-mortem from a male infant with SGBS ^30^ and can be differentiated *in vitro* to mature adipocytes. Of note, we observed higher activation of classic rather than 11-oxygenated androgens in SGBS cells (**Fig 3C**); incubations with 11KA4 yielded ample conversion to 11OHA4, with only a minor fraction of 11KA4 activated by AKR1C3 to 11KT and subsequently 11OHT (**Extended Data Fig. 2E**). These findings were entirely at odds with the pattern of androgen conversion we observed in the human female adipose tissue explant incubations, suggesting that SGBS cells do not represent a suitable model for studying human adipose tissue androgen metabolism.

### AKR1C3 inhibition selectively decreases 11-oxygenated androgen activation in human female adipose tissue

We first confirmed potent AKR1C3 inhibition by the selective AKR1C3 inhibitor BAY2299242 (BAY2) in non-steroidogenic HEK293T cells transfected to express AKR1C3 (IC50 33.1 ± 6.8 nM; **Fig. 4A**). In HEK293 cells transfected to express AKR1C3 together with the androgen receptor (AR) and an AR promotor reporter system, BAY2 blunted AR transactivation when the cells were treated with either A4 (**Fig. 4B**) or 11KA4 (**Fig. 4C**). Liquid chromatography-tandem mass spectrometry (LC-MS/MS) analysis of the supernatants from these experiments confirmed efficient inhibition of AKR1C3 activity (**Extended Data Fig. 3**).

**Fig. 4.**
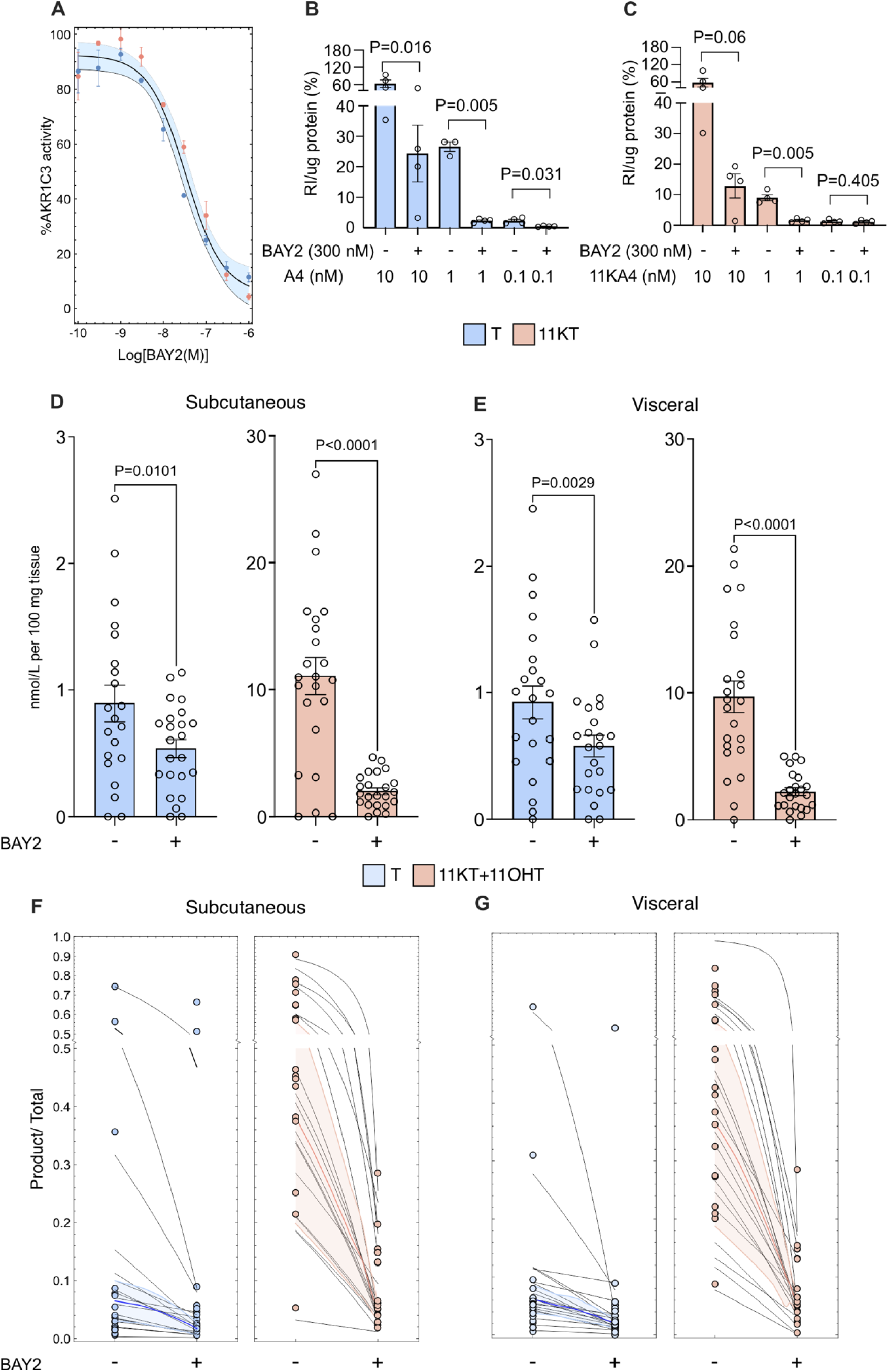
Effect of *in vitro* and *ex vivo* AKR1C3 inhibition on androgen and AR activation. (**A**) Inhibitor dose-response curves of AKR1C3 activity for the A4 to T and 11KA4 to 11KT reactions. HEK293T cells were transfected to express AKR1C3 and treated with either A4 or 11KA4 (100 nM) and increasing concentration of the AKR1C3 inhibitor BAY2299242 (BAY2) (n=3). Mean values with SEM as percentage of uninhibited activity are shown. A non-linear fit with the Four-Parameter Logistic (4PL) regression model with the slope parameter set to 1 was made to the model and the sigmoidal curve displays the 100% + response value with a 95% confidence interval. (**B & C**) A luciferase reporter assay in HEK293 cells transfected to constitutively express AKR1C3 and the AR, as well as luciferase under the control of an androgen response element, shows that AKR1C3 inhibition with BAY2 blocks AR transactivation in response to A4 and 11KA4 (n=4). Mean and SEM are shown. P-values were determined with a paired, two-tailed t-test. (**D & E**) Selective inhibition of 11-oxygenated androgen activation with BAY2 (300 nM) in female subcutaneous or visceral adipose tissue (n=22 donors, age 36-63, BMI 20.8-38.2 kg m^-2^) incubated *ex vivo* with androstenedione or 11KA4 (100 nM). Bars show the mean of all independent replicates ± SEM; circles represent the individual tissue donors. P-values were determined using Wilcoxon matched-pairs signed rank test. (**F & G**) *In silico* prediction of AKR1C3 inhibition in adipose tissue shows. A computational model of adipose tissue androgen metabolism by AKR1C3 and HSD11B1 was fitted to the experimental conversion data from the *ex vivo* subcutaneous (**F**) and visceral (**G**) adipose experiments from D & E in the absence of the AKR1C3 inhibitor (n=22). The reduction in AKR1C3 activity up to its fitted level of inhibition was then simulated and compared to the inhibition achieved in the *ex vivo* experiments with 300 nM BAY2 for each individual donor. The experimental data is shown as individual data points (open circles), while the model’s simulations of increasing AKR1C3 inhibition are shown by the connecting black lines. The colored lines and shaded areas show the median and median absolute deviation of the model’s predictions. The data is shown as the ratio of AKR1C3 substrates over total classic (A4 / (A4 + T) or 11-oxygenated androgens (11KT + 11OHT) / (11OHA4 + 11KA4 + 11KT + 11OHT)).

Next, we evaluated the effect of BAY2 on classic and 11-oxygenated androgen activation in female adipose tissue. We carried out *ex vivo* incubations with BAY2 and either of the two AKR1C3 substrates, A4 or 11KA4, in human female subcutaneous and visceral adipose tissue from 22 women undergoing elective gynecological surgery (age range 36-74 years, BMI range 20.8-38.2 kg m^-2^). This demonstrated a major decrease in the generation of active 11-oxygenated androgens (11KT and 11OHT) from 11KA4 in the presence of 300 nM BAY2, with only a minor reduction of T formation from A4 (**Fig. 4D and E**).

To further interrogate the results from our *ex vivo* experiments, we fitted a previously validated computational model to the *ex vivo* data to accurately predict the degree of AKR1C3 inhibition. This model ^28,31^ incorporates enzyme kinetic parameters for multiple enzymes involved in peripheral steroid metabolism, including AKR1C3 and HSD11B1, determined through *in vitro* experiments. Modelling of the *ex vivo* experimental data revealed that AKR1C3 inhibition primarily decreases 11-oxygenated androgen activation (**Fig. 4F and G**). It also demonstrated the extent of AKR1C3 inhibition needed to fit the experimental data obtained in the adipose tissue explant incubations (12.4 ± 2%, and 10.5 ± 2% residual AKR1C3 activity for subcutaneous and visceral adipose, respectively). These numbers are in good agreement with the inhibition observed in the HEK293T cells transfected with AKR1C3 after incubation with 300 nM BAY2 (14.2 ± 2.76 % residual activity, **Fig. 4A**), indicating that the observed effects of BAY2 in adipose tissue are exclusively due to AKR1C3 inhibition.

### AKR1C3 inhibitor treatment selectively decreases *in vivo* 11-oxygenated androgen activation in premenopausal women

Despite the ability of AKR1C3 to activate both 11-oxygenated and classic androgens (**Fig. 5A**), we demonstrated *ex vivo* and *in silico* that AKR1C3 inhibition selectively inhibited 11-oxygenated androgen activation in adipose tissue. We next performed comprehensive serum androgen profiling by LC-MS/MS in 21 premenopausal women treated with the selective AKR1C3 inhibitor BAY1128688 (BAY1) at a dose of 60 mg twice daily (n=9) or placebo (n=12) for 12 weeks in a phase 2a clinical trial ^32^. While we observed only a very minor effect of the AKR1C3 inhibitor on serum testosterone concentrations (**Fig. 5B**), results demonstrated a clear decrease in serum 11KT and 11OHT, both following single dose exposure at baseline and after 12 weeks of sustained once daily AKR1C3 inhibitor treatment (**Fig. 5 C and D**). There were no changes in circulating levels of the 11-oxygenated androgen precursors 11OHA4 and 11KA4 (**Extended Data Fig. 4B and C and Extended Data Fig 5B and C**). After the end of AKR1C3 inhibitor treatment, circulating 11OHT and 11KT concentrations increased back to baseline levels (**Fig. 5F and G**).

**Fig. 5.**
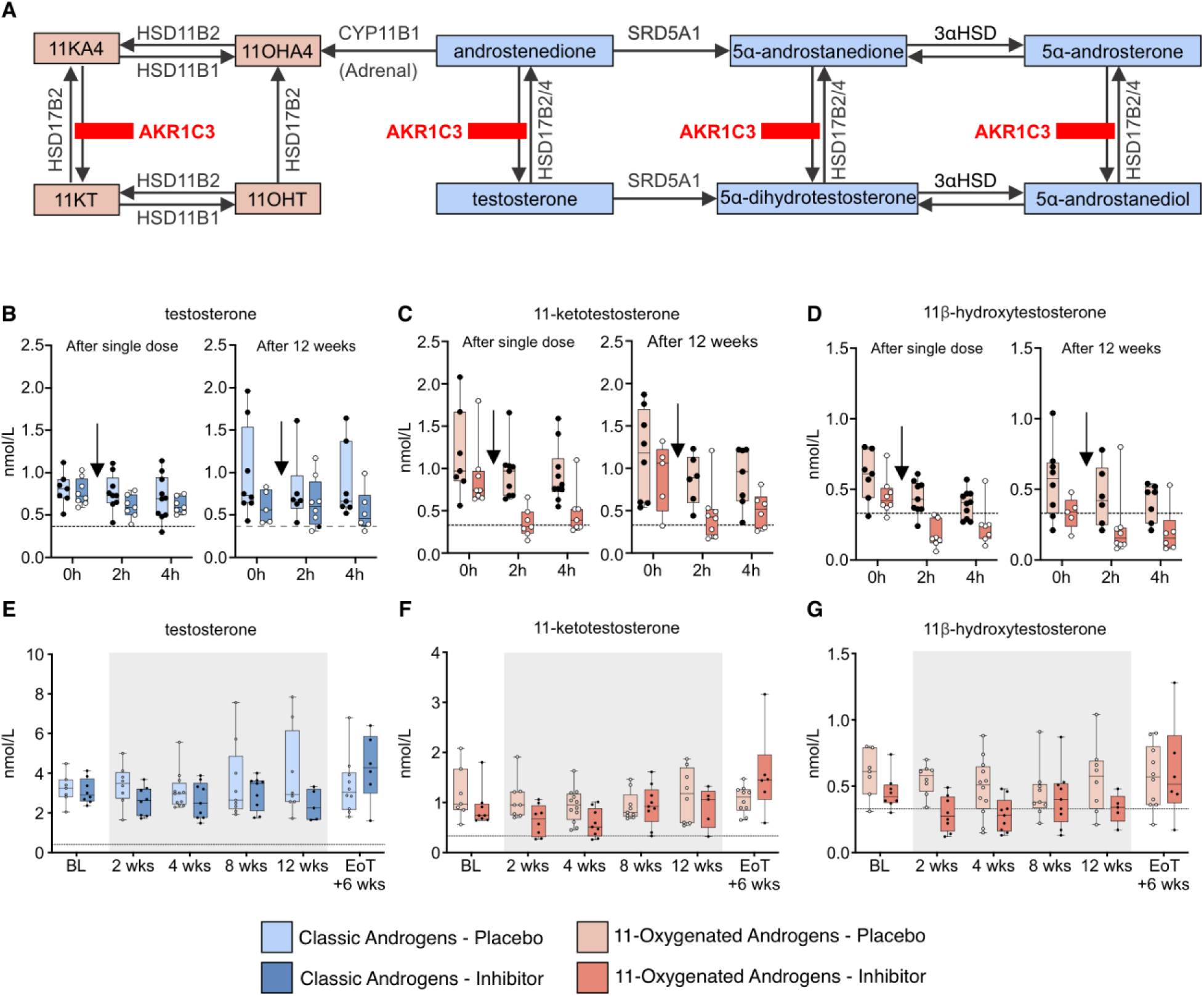
Serum androgen response to *in vivo* AKR1C3 inhibition. (**A**) Schematic showing the AKR1C3-catalyzed conversions for 11-oxygenated androgens (red) and classic androgens (blue). Serum concentrations of testosterone, 11-ketotestosterone and 11β-hydroxytestosterone over the 12 weeks of treatment with either 60 mg BAY1128688 (BAY1) taken orally twice daily (n=9) or placebo (n=12). Panels **B-D** show the acute androgen response to BAY1 and placebo within 4 hours of oral intake of inhibitor or placebo at baseline (=first dose) and after 12 weeks of treatment with no effect on testosterone but decreases in the circulating concentrations of 11KT and 11OHT at 2 and 4 hours. Panels **E-G** show the morning serum concentrations of active androgens during the 12-week treatment period and 6 weeks after the end of treatment (=EoT) with the AKR1C3 inhibitor or placebo. Serum steroid levels were quantified by LC-MS/MS. The arrow indicates the timing of inhibitor or placebo administration. Boxes show the median and 25^th^-75^th^ percentile, whiskers the range. Circles represent the individual participants. The dotted line indicates the lower limit of quantification for the respective analyte.

Over the treatment period of 12 weeks, alongside the decrease in 11-oxygenated androgens, we observed a progressive increase in morning serum levels of 5α-reduced steroids in the AKR1C3 inhibitor-treated premenopausal women. This included increases in the circulating concentrations of the potent androgen 5α-dihydrotestosterone (DHT) and the androgen metabolites 5α-androstanedione and 5α-androsterone (**Extended Data Fig. 6C-E**). This is suggestive of an upregulation of androgen activation via steroid 5α-reductase activity, which converts T to DHT. Circulating levels of 5α-reduced androgenic steroids returned to baseline within six weeks after the end of treatment (**Extended Data Fig. 6**). Of note, this increase in 5α-reduced steroids was not observed upon acute exposure to the AKR1C3 inhibitor, i.e. two and four hours after oral administration of BAY1, neither at baseline nor after 12 weeks of treatment (**Extended Data Fig. 4H to J**).

### Computational modelling indicates increased systemic AKR1C3 activity in women with polycystic ovary syndrome and predicts preferential inhibition of 11KT over T activation

We fitted the above-described computational model for the accurate prediction of AKR1C3 activity to serum steroid concentrations we measured in 91 treatment naïve women with PCOS (median age 28 years, range 19-48 years) and 66 women without PCOS of comparable age (median age 30 years, range 22-48 years; **Extended Data Table 2**). The computational model predicted increased systemic AKR1C3 activity in the PCOS cohort compared to controls (median 2.6-fold increase; **Fig. 6A**).

**Figure 6:**
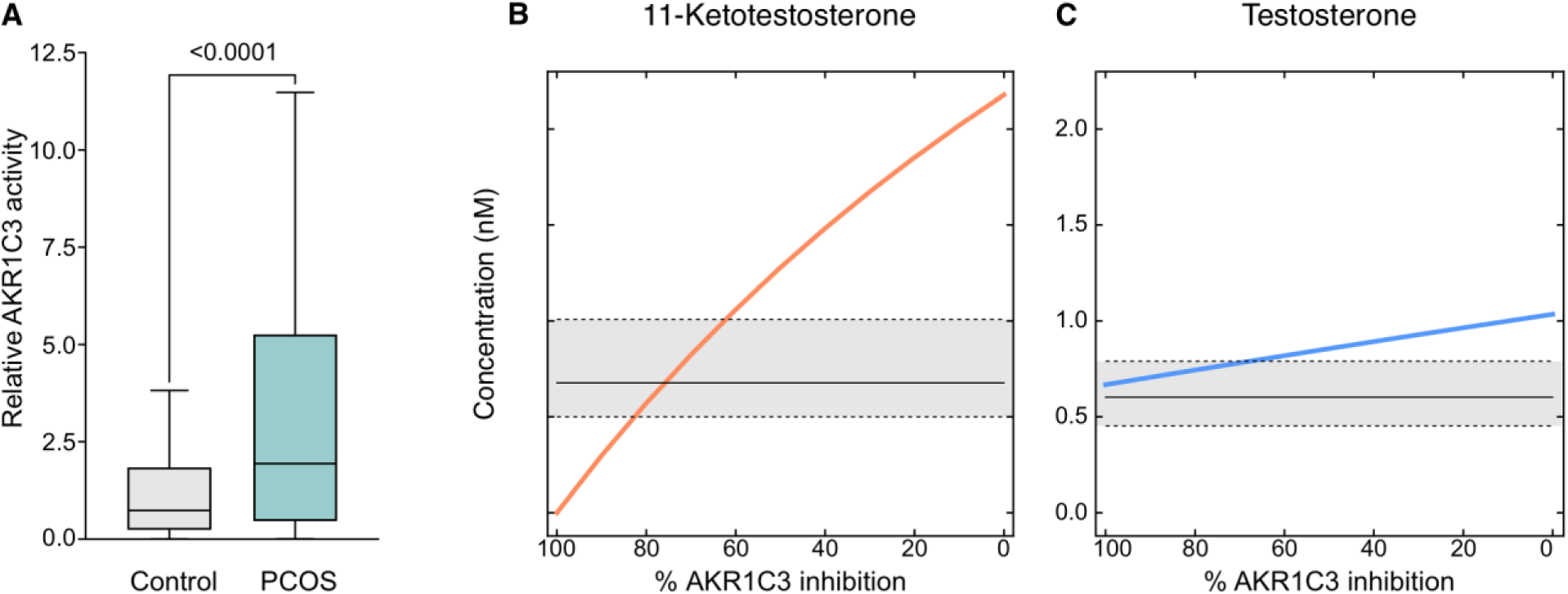
*In silico* prediction of systemic AKR1C3 activity and effect of AKR1C3 inhibition in women with PCOS. A computational model of peripheral glucocorticoid and androgen metabolism was fitted to the androgen and glucocorticoid serum profiles of treatment naïve women with women with PCOS (n=91) and an age-matched female control group (n=66). (**A**) The model was used to predict the relative AKR1C3 in the PCOS cohort in comparison to the healthy controls. Boxes show the median and 25^th^-75^th^ percentile, whiskers the range. P-values were determined using a Mann-Whitney U Test. The inhibition of AKR1C3 from 0 to 100% was then simulated and the effect on systemic T (**B**) and 11KT (**C**) predicted. The initial concentrations (0% inhibition) for T and 11KT are representative of the median concentration in the PCOS group as measured by LC-MS/MS. Solid horizontal lines denote the median concentrations, grey shaded areas the interquartile range (1^st^ to 3^rd^ quartiles) and dotted lines the 25^th^ and 75^th^ centile, respectively, of the sex- and age-matched control group.

Subsequent simulations of AKR1C3 inhibition from 0 to 100% in the PCOS group demonstrated that AKR1C3 inhibition resulted in higher reduction of 11-oxygenated over classic androgen activation (**Fig. 6B and C**). In addition, the model predicted that PCOS-related 11KT excess could be decreased to the median value of healthy women if AKR1C3 activity was reduced to 24% of uninhibited activity, while inhibition to 38% and 17% of residual activity decreased 11KT to the 75^th^ and the 25^th^ centile of controls, respectively (**Fig. 6B**). By contrast, even full inhibition of AKR1C3 did not decrease circulating T in PCOS to the median of controls; a decrease to the 75^th^ centile of T in controls could be achieved by inhibition to 32% of uninhibited activity (**Fig. 6C**).

## DISCUSSION

Androgen excess is a defining feature of PCOS and considered a key driver of metabolic dysfunction in affected women ^1,3–11^. Both the classic androgens, which originate from adrenal and gonadal biosynthesis, and the exclusively adrenal-derived 11-oxygenated androgens can be activated by the enzyme AKR1C3 ^16,33^ in peripheral target tissues of androgen action. A previous study in a healthy reference cohort reported an increase in circulating 11-oxygenated androgens with increasing BMI, pointing to adipose tissue as an important source; by contrast, BMI correlated negatively with circulating classic androgens ^21^. Previous studies have highlighted 11-oxygenated androgen excess in women with congenital adrenal hyperplasia ^34–36^, PCOS ^25^ and in individuals affected by one of the forerunner conditions of PCOS, premature adrenarche ^37^. Currently, androgen excess in PCOS is routinely assessed by measurement of serum T only ^1^. However, comprehensive androgen profiling using tandem mass spectrometry is starting to reveal that androgen excess in PCOS is heterogeneous, with a considerable proportion of women presenting with increased circulating 11-oxygenated androgens but normal testosterone ^25,26,38^, suggesting opportunities for a precision medicine approach based on the androgen profile. Here, we investigated the role of AKR1C3 in systemic and adipose-specific androgen homeostasis and the impact of AKR1C3 inhibition on circulating classic and 11-oxygenated androgens in women.

Using snRNA-Seq, we show that adipocytes are the major site of *AKR1C3* and *AR* expression within adipose tissue. *AKR1C3* expression levels were higher in subcutaneous than visceral adipose tissue, in agreement with previous studies using semi-quantitative RT-PCR ^13,28,39^. *AKR1C3* expression levels were higher in women than men and in obese than lean individuals, a distribution mirrored by the expression of *AR* in subcutaneous female adipose tissue. Using bulk RNA-Seq, we show that *AKR1C3* expression positively correlated with BMI in subcutaneous female but not male adipose tissue. Collectively, these data demonstrate major sex differences in the expression of genes crucial for mediating biological androgen action, indicating that adipose tissue is a key site of peripheral androgen activation and action in women. Previous work has shown AKR1C3-mediated androgen activation results in lipid accumulation in the adipocyte and promotes a systemic lipotoxic metabolome ^12,14^. This in turn results in enhanced insulin secretion, which drives increased *AKR1C3* expression in adipocytes ^12,15^, thereby creating a vicious circle linking AKR1C3-mediated androgen activation with increased insulin resistance ^12^. This highlights AKR1C3 as a potential pharmaceutical target for treating androgen excess and the accompanying metabolic dysfunction in women with PCOS, who regularly exhibit both androgen excess and insulin resistance.

Previous work based on *in vitro* biochemical experiments reported higher catalytic efficiency of AKR1C3 towards 11-oxygenated androgen precursors ^15,40^. In this study, using *ex vivo* incubations of subcutaneous and visceral adipose tissue explants, we found that female adipose tissue preferentially activates 11-oxygenated over classic androgens. When incubating the human female adipose tissue explants with a selective small molecule AKR1C3 inhibitor (BAY2299242, BAY2), we observed a major decrease in 11-oxygenated androgen activation, with only a minor impact on classic androgen activation. Excitingly, we observed the same pattern of selective disruption of 11-oxygenated over classic androgen activation in our 12-week *in vivo* study with another, chemically distinct AKR1C3 inhibitor (BAY1128688, BAY1; steroid backbone structure), both after single dose exposure and sustained treatment for 12 weeks.

It is important to consider that steroid metabolism and the effects of its manipulation should not be looked at in isolation, i.e. by looking at a single enzyme only. When treating premenopausal women with AKR1C3 inhibitor for 12 weeks, we observed over time an increase in systemic 5α-reductase activity as indicated by increasing circulating concentrations of 5α-reduced steroids including the potent androgen 5α−DHT. Further studies will need to assess the clinical significance of these findings in comparison to the pronounced reduction in 11-oxygenated androgens, the androgen class we have shown here to be primarily activated physiologically within female adipose tissue. However, a recent study demonstrated that adipose tissue is protected from the activity of systemic 5α-reduced androgens due to its high expression of AKR1C2, which readily inactivates 5α-reduced androgens ^41^. Moreover, all women in our *in vivo* study had normal or low circulating androgens, which were decreased even further by AKR1C3 inhibition. Therefore, it is readily conceivable that the systemic upregulation of 5α-reductase activity, observed in this 12-week phase 2a study, is a counter-regulatory response to androgen deficiency induced by AKR1C3 inhibitor treatment. It is in no way certain that an upregulation of systemic 5α-reductase activity would also be observed when administering AKR1C3 inhibitors to women with PCOS-related androgen excess where the goal would be the normalization of pathologically increased androgen levels.

In our adipose tissue studies, we also show that the enzyme HSD11B1 is co-expressed with AKR1C3 in adipocytes, albeit at much lower levels. While HSD11B1 is well known as a glucocorticoid-activating enzyme, it can also inactivate 11-oxygenated androgens, by converting 11KT to the less active 11OHT ^28^. Conversely, HSD11B1 has recently been shown to activate classic androgens ^29^ and it is possible that the persistence of classic androgen activation in adipose tissue despite AKR1C3 inhibition is driven by HSD11B1 activity. However, the by far higher activation rate of 11-oxygenated over classic androgens we observed in adipose tissue will be primarily driven by the substrate preference of AKR1C3 for 11-oxygenated androgen precursors, corroborating previous *in vitro* findings ^40^.

Strengths of our study include its integrated translational approach, featuring complementary *in vitro*, *ex vivo*, *in vivo* and *in silico* experiments to comprehensively investigate the effect of AKR1C3 inhibition on androgen activation in women, both at tissue-specific and systemic levels. A limitation is that our *ex vivo* experiments focused only on adipose tissue while other sites of AKR1C3 expression may also contribute to systemic 11KT levels. In addition, it remains to be established how AKR1C3 inhibition and the resulting reduction in 11-oxygenated androgen activation impact upon adipose biology and metabolic function.

Taken together, we show that AKR1C3 inhibition selectively decreases the activation of 11-oxygenated androgens, both systemically and within adipose tissue. Applying our validated computational model ^28,31^ to the measured serum androgen profiles, we also show that systemic AKR1C3 activity is increased in women with PCOS. Thus, the specific targeting of 11-oxygenated androgen activation by AKR1C3 inhibition can be considered an important step towards a precision medicine approach to the treatment of women with PCOS. Further taking into account the reported link of AKR1C3 to insulin resistance ^12,14^, we propose that AKR1C3 inhibition represents not only a highly promising therapeutic approach to PCOS-related androgen excess but may also hold potential for ameliorating PCOS-related metabolic dysfunction, which warrants exploration by future studies.

## MATERIALS AND METHODS

### Experimental design

Gene expression across various human tissues was studied in a sex-specific analysis in publicly available bulk RNA-Seq data (The Adult Genotype Tissue Expression (GTEx) Project) and in a snRNA-Seq data sets from subcutaneous adipose tissue of 49 donors ^27^. Experiments with cell lines and human tissue were designed to study the effect of pharmacological AKR1C3 inhibition on the generation of active androgens. For cell line models at least three biologically independent experiments were performed. Due to intra-individual variation in AKR1C3 expression, experiments using human tissue were performed including tissue from at least eight donors. To study the effect of AKR1C3 inhibition on androgen levels *in vivo*, we analyzed serum samples from a phase 2a clinical trial with selective AKR1C3 inhibitors. We also carried out comprehensive androgen profiling in women with PCOS in comparison to healthy female controls of comparable age. Due to the exploratory nature of our analysis, participant numbers were predetermined by the availability of samples from these clinical studies. All tissue donors and clinical study participants were female. Experimental and clinical studies were complemented by a computational model of peripheral steroid metabolism, which was fitted to circulating steroid concentrations.

### Gene expression analysis

Gene expression data as transcript per million (TPM) and body mass index of the participants were downloaded from The Adult Genotype Tissue Expression (GTEx) Project database (dbGaP Accession phs000424.v10.p2; n=233 for female subcutaneous adipose tissue, n=481 for male subcutaneous adipose tissue, n=108 for female visceral adipose tissue and n=407 for male visceral adipose tissue). Descriptive statistics and correlation analysis were performed in R using ggplot.

The subcutaneous adipose tissue snRNA-seq atlas was established as previously described^27^ combining data from 34 (25 female) individuals with obesity (BMI > 30 kg/m^2^) and 28 (18 female) lean controls (BMI < 26kg/m^2^) ^27,42^.

### Reagents and plasmids

11β-hydroxyandrostenedione (11OHA4), 11-ketoandrostenedione (11KA4), androstenedione (A4) and testosterone (T) were purchased from Sigma-Aldrich, while 11β-hydroxytestosterone (11OHT) and 11-ketotestosterone (11KT) were obtained from Steraloids. The AKR1C3 inhibitor BAY2299242 (BAY2) was provided by BAYER AG, Berlin, Germany. Stock solutions of the steroids (1 mg/mL in methanol) and inhibitors (10 mM in DMSO) were stored at -80 °C.

pcDNA3 with human AKR1C3 was obtained from Prof Jerzy Adamski (Helmholtz Zentrum München, Germany). pcDNA3 with the human androgen receptor was obtained from Prof Jeremy W Tomlinson (University of Oxford, UK). pTAT-GRE-E1b-luc was obtained from Prof Guido Jenster (Erasmus University of Rotterdam, Netherlands). pCMV8 was purchased from the American Type Culture Collection ATCC.

### Adipose tissue *ex vivo* androgen metabolism assay

Adipose tissue samples were transported to the laboratory at room temperature in Dulbecco’s Modified Eagle’s Medium/Nutrient Mixture F-12 Ham (Sigma-Aldrich) supplemented with 100 U/mL penicillin, 0.1 mg/mL streptomycin, 33 µM biotin and 17 µM panthotenic acid (Sigma-Aldrich). Connective tissue and vessels were removed and the adipose tissue washed in phosphate buffered saline. Tissue samples of 100-300 mg were cut and weighed. Each sample was cut into four smaller pieces for incubations with 100 nM steroid with AKR1C3 inhibitor or vehicle in media supplemented as described above. Tissue incubations were constantly rotated at 37 °C for 24 or 72 hours. The supernatant of the culture medium after centrifugation (10 minutes at 16,000*g* and 4 °C) was snap frozen in liquid nitrogen and stored at -80 °C.

### Androgen metabolism assay using Simpson-Golabi-Behmel syndrome adipocytes

The human Simpson-Golabi-Behmel syndrome (SGBS) preadipocyte cell line was proliferated in DMEM/Nutrient Mixture F12 Ham (DMEM/F12) supplemented with 10% fetal bovine serum, 33 μM biotin, 17 μM pantothenate, 100 U/mL penicillin and 0.1 mg/mL streptomycin at 37 °C in 90% humidity and 5% CO2 for a maximum of 52 generations to ensure high capacity for differentiation ^30^. SGBS preadipocytes were differentiated into lipid-storing adipocytes in serum-free DMEM/F12 supplemented with 100 U/mL penicillin, 0.1 mg/mL streptomycin, 33 μM biotin, 17 μM pantothenate, 0.01 mg/mL transferrin, 20 nM insulin, 100 nM cortisol, 0.2 nM triiodothyronine, 25 nM dexamethasone, 2 μM rosiglitazone and 500 μM methyl-3-isobutylxanthine for 10-11 days according to established protocols ^30^. After differentiation, SGBS cells were treated with serum-free DMEM/F12 (33 μM biotin, 17 μM pantothenate and 1% penicillin-streptomycin) containing 100 nM steroid substrate plus 300 nM AKR1C3 inhibitor or vehicle. After 24 hours, the medium was collected for analysis.

### Steroid quantification by LC-MS/MS

200 µL of serum or medium from cell line or tissue incubations were extracted by liquid-liquid extraction and steroids were quantified by LC-MS/MS using a validated assay ^21,43^.

### AKR1C3 activity assay in HEK293T cells

HEK293T cells were purchased from ATCC and cultured in Dulbecco′s Modified Eagle′s Medium (high glucose, Sigma-Aldrich) supplemented with 10% fetal bovine serum (Sigma Aldrich), 100 U/mL penicillin and 0.1 mg/mL streptomycin (Gibco) at 37 °C, 90% humidity and 5% CO2. Cells were transfected with pcDNA3-AKR1C3 (17 μg) using FuGENE® HD transfection reagent (Promega) in 100-mm dishes. 24 hours after transfection cells were plated in 96-well plates at 1.5x10^4^ cells/well. 24 hours later the culture medium was replaced by 200 μL of serum-free medium containing 100 nM A4 or 11KA4 with increasing concentrations of the AKR1C3 inhibitors. Medium samples were collected after four hours for 11KA4 and six hours for A4 for steroid quantification by LC-MS/MS. For the determination of IC50 values, mean values of the percentage product formation compared to the vehicle treated samples were plotted against the inhibitor concentration and IC50 were determined using a non-linear fit.

### AR transactivation luciferase reporter assay

Androgen receptor transactivation was studied in HEK293 cells (ATCC) transiently transfected with AKR1C3 and a promoter luciferase reporter assay similar to previously described assays ^19,44^. HEK293 cells in 75-cm^2^ flasks were co-transfected with pcDNA-AKR1C3 (12 μg), pcDNA3-AR (2 μg) and pTAT-GRE-E1b-luc (10 μg; containing a glucocorticoid receptor response element, which is also recognized by AR, and the luciferase reporter construct under the control of the Eb1 promoter) using FuGENE® HD transfection reagent. Co-transfections with pcDNA-AKR1C3, empty pCMV8 and pTAT-GRE-E1b-luc were performed as controls to establish that observed effects are AR-mediated. 24 hours after transfection cells were plated in 24-well plates at 10^5^ cells/well in 500 μL culture medium. After a further 24 hours the culture medium was replaced by serum-free medium supplemented containing A4 or 11KA4 and 300 nM of the AKR1C3 inhibitor BAY2 or vehicle. After 24 hours the medium was removed for steroid quantification by LC-MS/MS and luciferase activity in the cell lysates was determined using the Luciferase Assay System (Promega, E1500). Luminescence was normalized to the total protein content of the respective well determined using the DC Protein Assay (Bio-Rad).

### Computational modelling of steroid metabolism and enzyme inhibition

A computational model of peripheral androgen and glucocorticoid metabolism was previously constructed using the parameterized rate equations (enzyme kinetics) for the enzymes AKR1C3, HSD17B2, HSD11B1 and HSD11B2 ^28,31^. Ordinary differential equations were defined for each individual steroid substrate and product in the model. The model was used to predict the relative activity levels of AKR1C3 from each individual adipose tissue donor used in the *ex vivo* adipose incubations by fitting the model to the experimentally determined steroid conversion data and minimizing the sum of the squared differences between the simulated and measured steroids levels. Similarly, the model was used to predict the relative systemic activity of AKR1C3 in each individual women with PCOS or healthy control by assuming steady state and minimizing the sum of the squared differences between the simulated and measured steroids levels. For the minimization, a multiplier for the individual enzymes was fitted relative to the activity of HSD11B2 which was set to 1. Following the estimation of relative enzyme activities, simulations of AKR1C3 inhibition were performed in silico. A full model description is available here: 10.6084/m9.figshare.27043891.

### Clinical cohorts and human biomaterial

Paired subcutaneous and visceral adipose tissue samples for use in the ex vivo incubations with and without AKR1C3 inhibitor were collected from women undergoing elective gynecological, non-cancer surgery (11oxo-PCOS; ClinicalTrials.gov ID NCT05246865; UK Integrated Research Assessment System (IRAS) number 221491; Research Ethics Committee (REC) reference 19/WM/0183) or bariatric surgery (ClinicalTrials.gov ID NCT02124486; IRAS REC reference 14/WM/0011). Written, informed consent was given by all participants prior to surgery and sample collection.

Serum samples for androgen metabolome profiling by tandem mass spectrometry were collected through two different studies: (1) a randomized, placebo-controlled, double-blind, multi-center phase 2a clinical trial with continuous daily administration of the AKR1C3 inhibitor BAY1128688 (BAY1) for 12 weeks in premenopausal women with endometriosis (AKRENDO; ClinicalTrials.gov ID NCT03373422; EudraCT Number 2017-000244-18); and (2) an observational study for detailed phenotyping of treatment-naïve women with polycystic ovary syndrome (DAISy-PCOS; ClinicalTrials.gov ID NCT03911297; IRAS number 256572, and REC reference 19/WM/0110). The healthy female controls were taken from a previously published reference range data set measured with the same assay ^21^.

### Statistical analysis

Gene expression was compared by Wilcoxon signed-rank test or Mann-Whitney U Test as appropriate. Correlation between BMI and gene expression was assessed using Pearsons’s correlation analysis. Analysis of the effect of AKR1C3 inhibition was performed using paired, two-tailed T-test for the AR transactivation assay, and by Wilcoxon matched-pairs signed rank test for the androgen metabolism in tissue *ex vivo*.

## Acknowledgements

**Funding:**

Wellcome Trust Investigator Grant WT209492/Z/17/Z (WA) Medical Research Council UK grant MC_UP_1605/15 (WA)

Academy of Medical Sciences UK Newton Advanced Fellowship NAF004\1002 (KHS) National Research Foundation of South Africa Competitive Programme for Rated Researchers SRUG2204052159 (KHS)

Wellcome Trust Clinical Career Development Fellowship 219602/Z/19/Z (WRS) Medical Research Council UK grant MR/K002414/1 (WRS)

Medical Research Council UK grant MC_UP_1605/7 (WRS)

National Institute for Health Research Imperial Biomedical Research Centre grant CL-2018-21-501 (WRS)

Diabetes UK grant 22/0006436 (WRS)

Health Research Board Emerging Clinician Scientist Award ECSA-2020-001 (MWO’R)

South African Research Chairs Initiative by the Department of Science and Technology and the National Research Foundation of South Africa grant 82813 (JLS)

Bayer AG (MP, AW, JPI, JH)

German Federal Ministry of Research, Technology and Space (Bundesministerium für Forschung, Technologie und Raumfahrt, BMFTR) as part of the German Center for Child and Adolescent Health (DZKJ) grant 01GL2407A (MW)

## Author contributions

Conceptualization: WA

Methodology: LS, LBLW, AET, MW, MWO’R, MP, AW, JPI, JLS, WRS, JH, KHS, WA

Investigation: LS, AVA, LBLW, AET, IO, AM, EM, TM, PK, PS, TJC, WRS, JLS, JH, KHS, WA

Visualization: LS, AVA, LBLW, JLS, KHS, WA

Funding acquisition: JLS, MWOR, WRS, KHS, WA

Project administration: WA

Supervision: WRS, JH, KHS, WA

Writing – original draft: LS, AVA, LBWL, KHS, WA

Writing – review & editing: all other authors

## Data and materials availability

All data associated with this study are presented in the paper or the Extended Data.

## Extended Data

**Extended Data Table 1:** Statistical significance of differences in *AKR1C3*, *HSD11B1* and androgen receptor (*AR*) expression in subcutaneous adipocytes from the snRNA-Seq atlas established with tissue from 34 (25 female) individuals with obesity (BMI > 30 kg/m2) and 28 (18 female) lean individuals (BMI < 26 kg/m^2^) ^27,42^. Expression was compared by Wilcoxon signed-rank test and Benjamini-Hochberg correction. FDR, false discovery rate.

| Gene | Comparison |  | p-value | FDR |
| --- | --- | --- | --- | --- |
| *AKR1C3* | lean vs obesity | female | 2.28e-207 | 4.51e-205 |
|  |  | male | 4.39e-91 | 1.34e-88 |
|  | male vs female |  | 1.76e-39 | 9.54e-38 |
| *HSD11B1* | lean vs obesity | female | 8.87e-117 | 7.67e-115 |
|  |  | male | 6.46e-24 | 3.85e-22 |
|  | male vs female |  | 0.008 | 0.036 |
| *AR* | lean vs obesity | female | 8.22e-148 | 9.30e-146 |
|  |  | male | 8.39e-11 | 2.51e-09 |
|  | male vs female |  | 6.71e-09 | 7.08e-08 |

**Extended Data Table 2:** Serum steroid concentrations measured by liquid chromatography-tandem mass spectrometry in 91 treatment-naïve women with PCOS and 66 age-matched female controls. All data represent medians and interquartile ranges.

|  | Women<br>with PCOS | Healthy female<br>controls |
| --- | --- | --- |
| Age (years) | 28 (25-32) | 30 (26-38.5) |
| DHEAS (μmol/L) | 5.8 (4.0-8.1) | 4.6 (3.4-6.0) |
| DHEA (nmol/L) | 20 (15-28) | 9.5 (6.5-14.0) |
| Androstenedione (A4) (nmol/L) | 6.1 (4.8-9.2) | 2.2 (1.6-3.5) |
| Testosterone (T) (nmol/L) | 1.5 (1.2-2.1) | 0.6 (0.5-0.8) |
| 5α-dihydrotestosterone (DHT) (nmol/L) | 0.43 (0.27-0.61) | 0.32 (0.22-0.48) |
| 11β-hydroxyandrostenedione (11OHA4) (nmol/L) | 8.9 (6.0-14) | 7.5 (5.4-10.3) |
| 11-ketoandrostenedione (11KA4) (nmol/L) | 3.4 (1.9-6.6) | 3.2 (2.3-3.9) |
| 11-ketotestosterone (11KT) (nmol/L) | 1.5 (1.1-2.1) | 0.6 (0.5-0.9) |
| 11β-hydroxytestosterone (11OHT) (nmol/L) | 0.78 (0.57-1.1) | 0.36 (0.25-0.51) |

**Extended Data Fig. 1.**
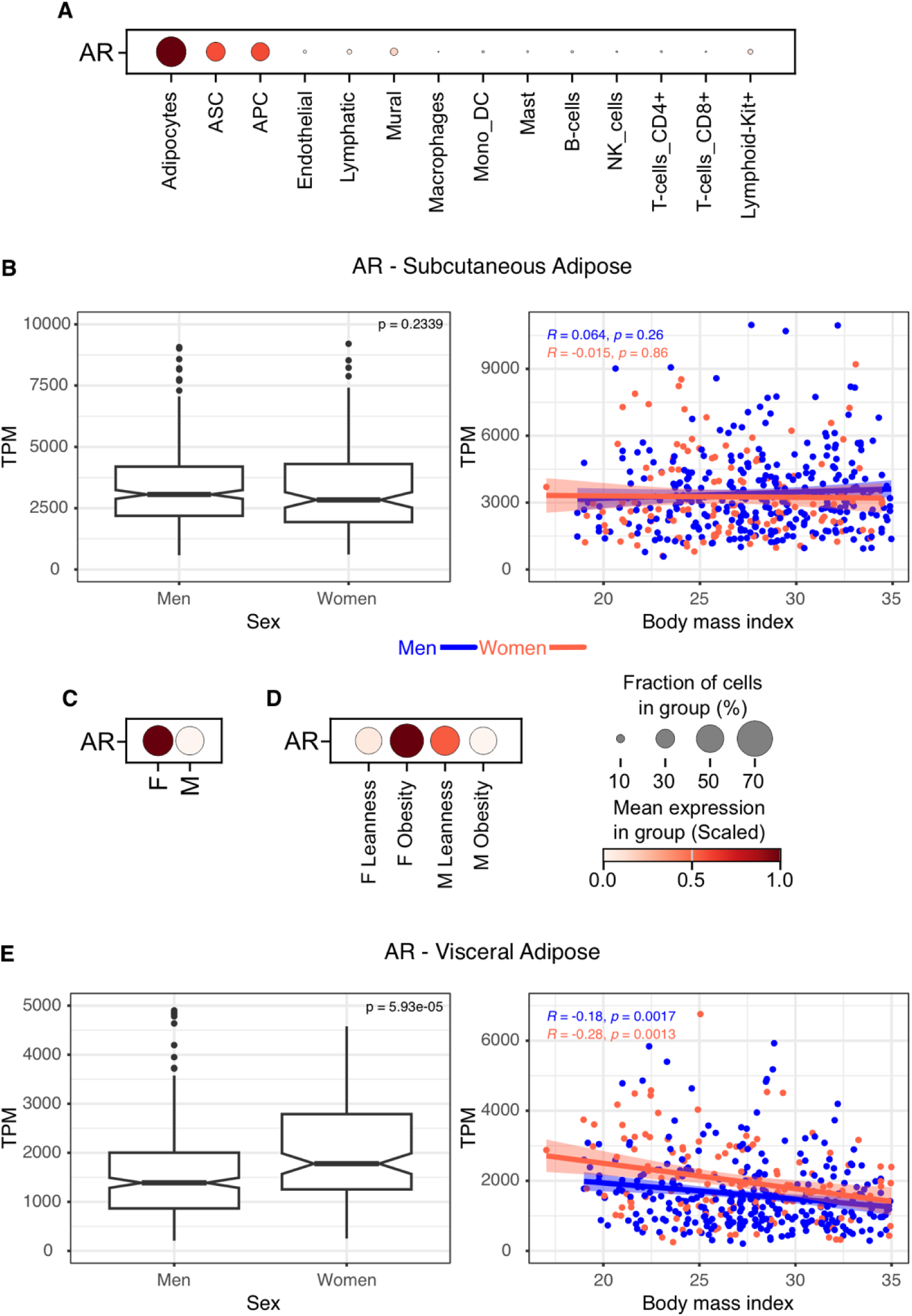
Expression of androgen receptor *(AR)* in human adipose tissue and the relationship with sex and body mass index (BMI). (**A**) Dot plot (mean expression scaled) from snRNA-Seq of human subcutaneous adipose tissue (n=49) showing that *AR* expression is most abundant in adipocytes (ASC, adipose-derived stem cells; APC, adipose progenitor cells; Mono_DC, monocyte-derived dendritic cells; NK_cells, natural killer cells). Analysis of bulk RNA-Seq data from the Adult Genotype-Tissue Expression (GTEx) Project (dbGaP Accession phs000424.v10.p2) showing the relationship of sex (Tukey box-and-whisker plots, p-values determined by Mann-Whitney U test) and BMI (Pearson correlation with 95% confidence interval) with *AR* expression in subcutaneous (**B**) and visceral adipose tissue (**C**). (**D**) Dot plot (mean expression scaled) from snRNA-Seq analysis of subcutaneous adipocytes from female (F, n=33) and male (M, n=16) donors showing the differential expression of *AR*. (**E**) Dot plot (mean expression scaled) from snRNA-Seq analysis of *AR* expression in subcutaneous adipose tissue from 24 individuals with leanness (15 female) and 25 with obesity (18 female).

**Extended Data Fig. 2.**
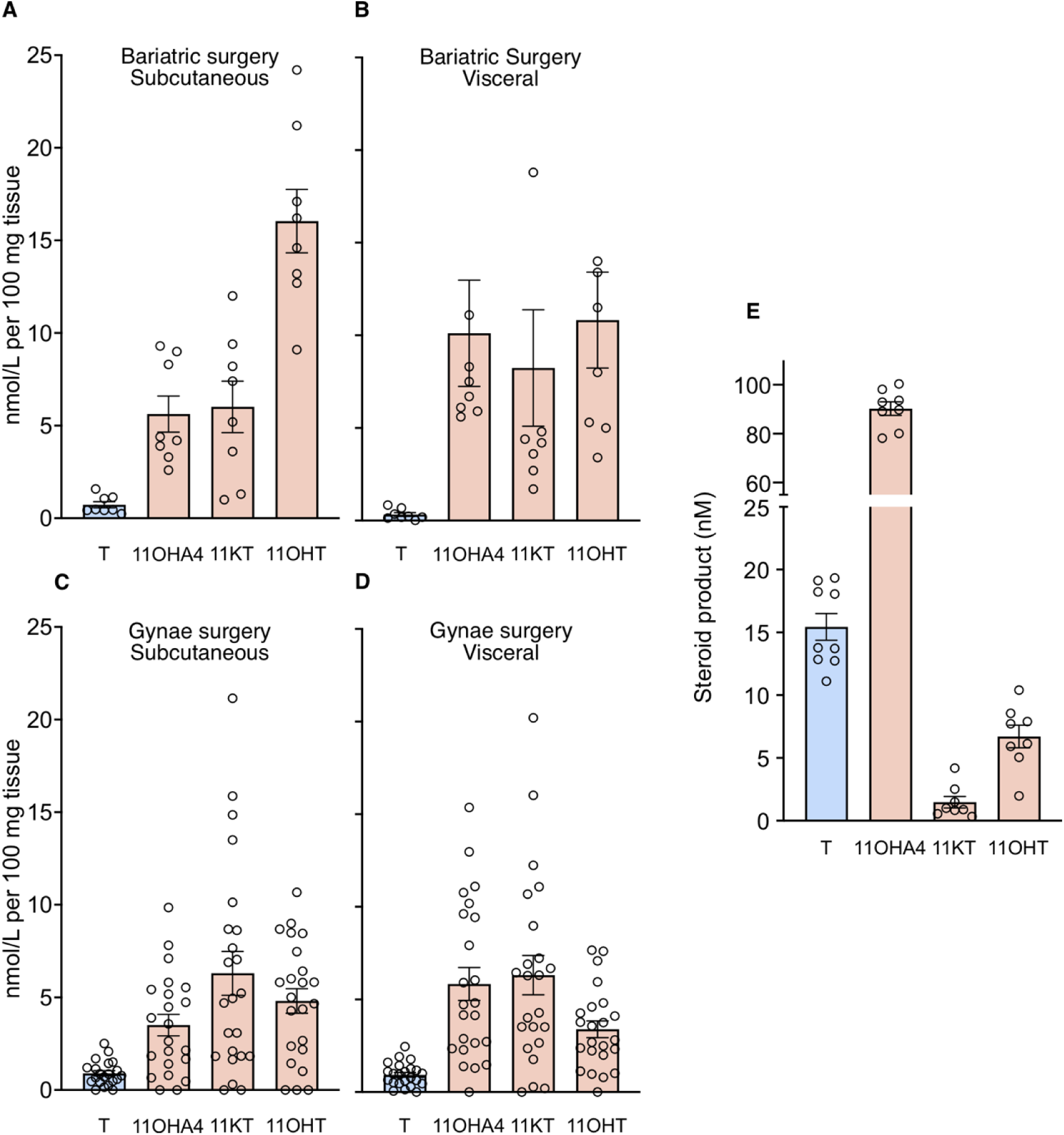
Comprehensive steroid profiles of classic and 11-oxygenated androgen metabolism in human female adipose tissue explants and SGBS cells (Fig 3). Incubations of subcutaneous and visceral adipose tissue showed preferential generation of active 11-oxygenated androgens (11KT and 11OHT) over the classic androgen (T). Adipose tissue was collected from women undergoing (**A & B**) bariatric surgery (n=8, age 32-59, BMI 42.0-56.8 kg/m^2^) and (**C & D**) gynecological surgery (n=22, age 36-63, BMI 20.8-38.2 kg/m^2^) and incubated with the AKR1C3 substrates androstenedione or 11KA4 (100 nM) for 72h (bariatric surgery) or 24h (gynae surgery). Androgen metabolites were quantified by LC-MS/MS. (**E**) Incubation of differentiated SGBS adipocytes with A4 or 11KA4 (100 nM) results in greater generation of the active classic androgen (T) compared to the 11-oxygenated androgens (11KT and 11OHT; n=8 or 9). Bars show the mean of all independent experiments ± IQR; circles represent the results in individual adipose tissue explants (**A-D**) or independent cell culture experiments (**E**).

**Extended Data Fig. 3.**
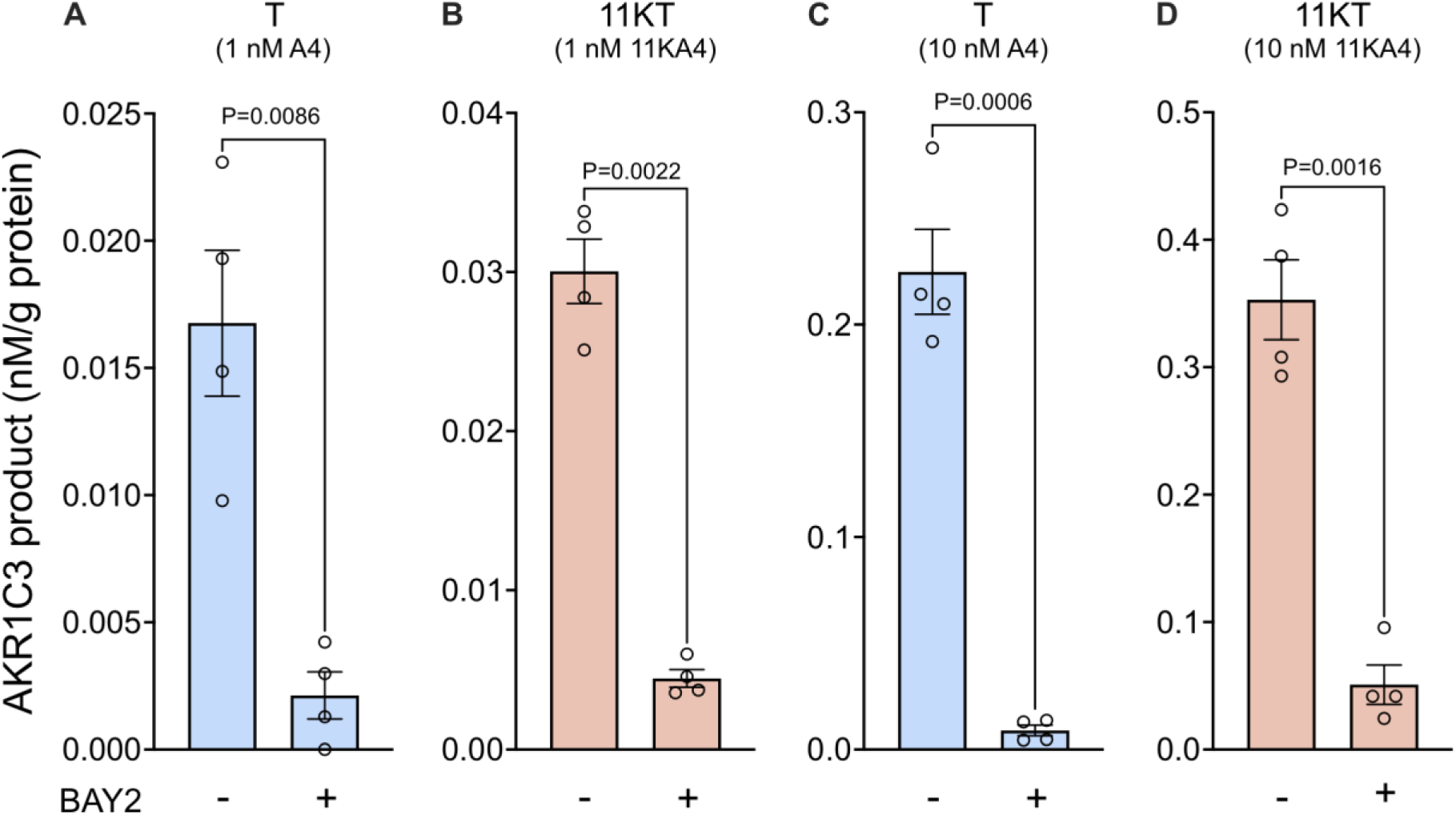
Effect of AKR1C3 inhibition on androgen activation in HEK293 reporter cells used in the AR transactivation assay (Fig. 4B and C). LC-MS/MS analysis demonstrates the inhibition of the generation of T (Panels **A & C;** incubations with 1nM and 10nM A4, respectively) and 11KT (Panels **B & D;** incubations with 1nM and 10nM 11KA4, respectively) in the presence of the AKR1C3 inhibitor BAY2, which supports the observed reduction in AR transactivation **(Fig. 4B and C)**. Bars show the mean of all independent experiments (n=4) ± SEM; circles represent the individual experimental values. P-values were determined using paired t-test.

**Extended Data Fig. 4.**
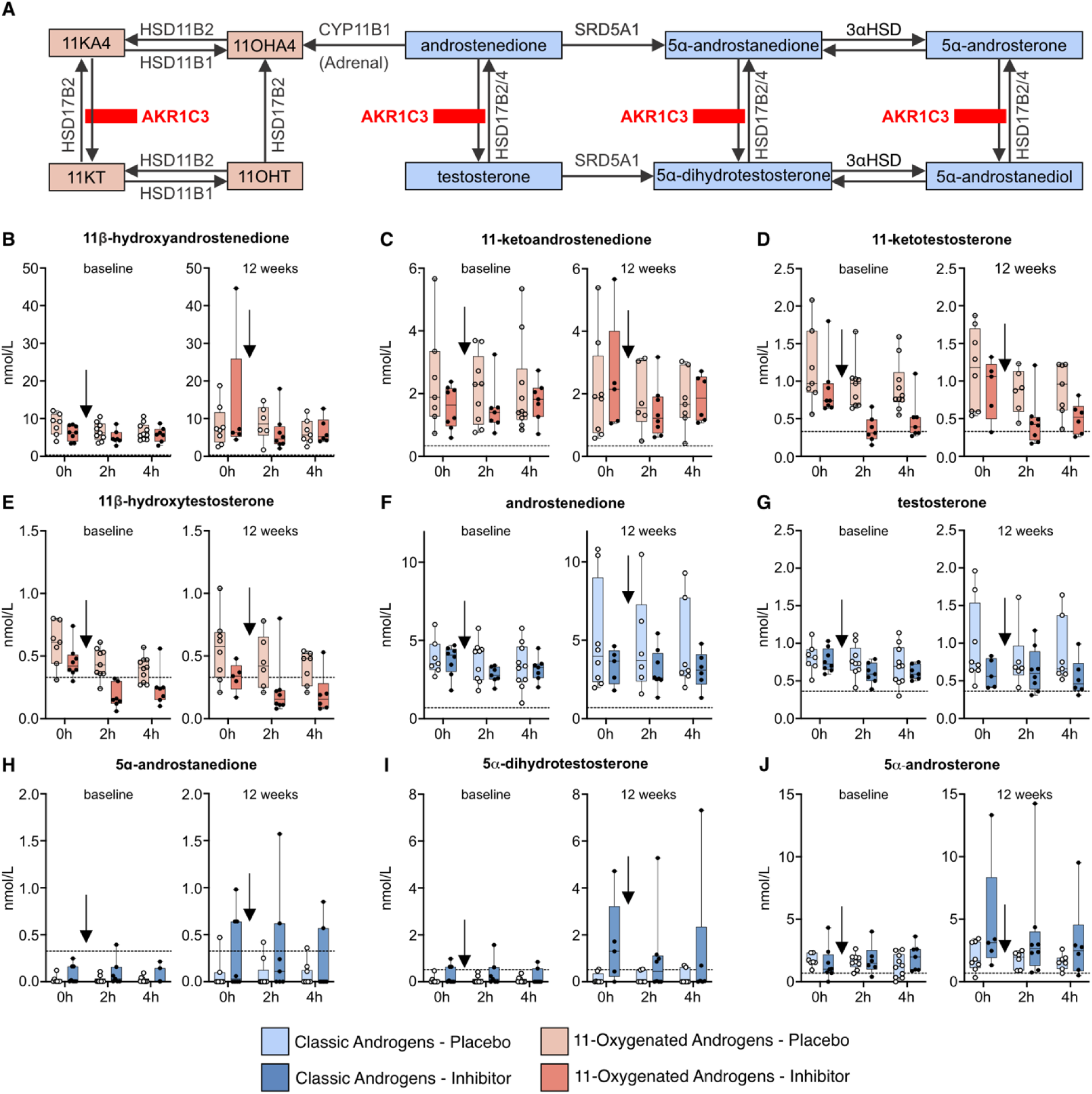
Acute serum steroid response to in vivo AKR1C3 inhibition in adult premenopausal women at baseline (=first dose) and after 12 weeks of treatment with BAY1128688 (BAY1). Women received either 60 mg BAY1 twice daily (n=9) or placebo (n=12). (A) Schematic showing the AKR1C3 catalyzed conversions for 11-oxygenated androgens (red) and classic androgens (blue). Serum levels of (B-E) 11-oxygenated androgens and (F-J) classic androgens quantified by LC-MS/MS show acute decreases in the active 11-oxygenated androgens (11KT and 11OHT, D & E) without discernible acute effect on classic androgens. The arrow indicates the timing of inhibitor or placebo administration. Boxes show the median and 25th-75th percentile, whiskers the range. Closed circles represent the individual participants. The dotted line indicates the lower limit of quantification of the LC-MS/MS assay.

**Extended Data Fig. 5.**
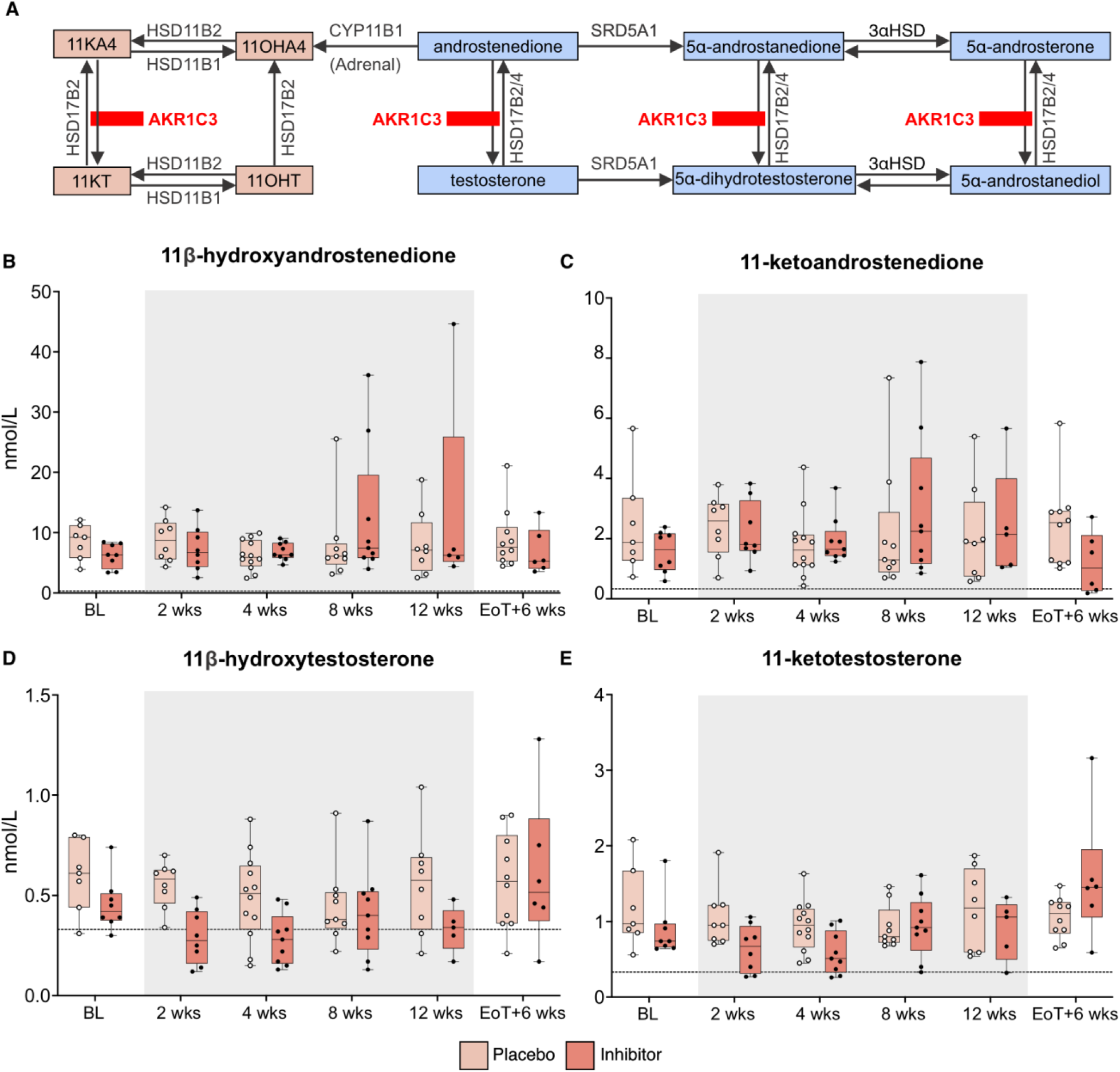
Effect of 12 weeks of treatment with the AKR1C3 inhibitor BAY1128688 (BAY1) on circulating 11-oxygenated androgens in premenopausal women. (**A**) Schematic showing the AKR1C3 catalyzed conversions for classic and 11-oxygenated androgens. (**B-E**) Serum steroids quantified by LC-MS/MS at study baseline (BL), during the 12-week treatment period (grey box) with either BAY1 (60mg bd; n=9) or placebo (n=12), and 6 weeks after the end of treatment (EoT), showing selective reductions of the AKR1C3 product 11KT and its metabolite 11OHT in individuals receiving the AKR1C3 inhibitor.

**Extended Data Fig. 6.**
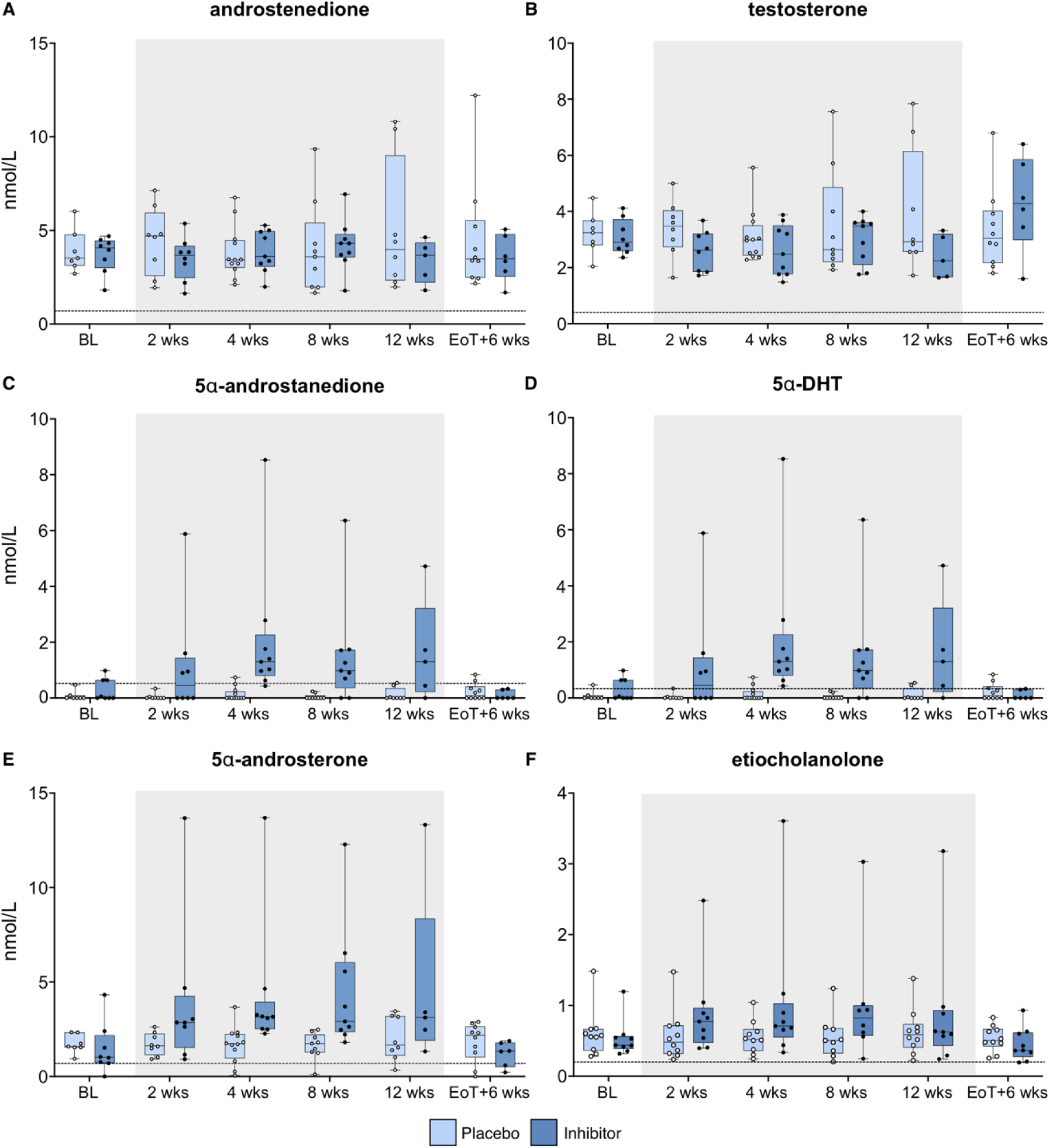
Effect of 12 weeks of treatment with the AKR1C3 inhibitor BAY1128688 (BAY1; 60 mg bd) on circulating classic androgens in premenopausal women. Serum steroid quantification by LC-MS/MS at study baseline (BL) and during the 12-week treatment period (grey box) with either BAY1 (60mg bd; n=9 or placebo (n=12) shows only a minor effect on serum androstenedione and testosterone (**A & B**), but an increase in 5α-reduced steroids during the treatment period, reverting to pre-treatment levels 6 weeks after the end of treatment (EoT).

